# The network of SARS-CoV-2—cancer molecular interactions and pathways

**DOI:** 10.1101/2022.04.04.487020

**Authors:** Pau Erola, Richard M. Martin, Tom R. Gaunt

## Abstract

**Background:** Relatively little is known about the long-term impacts of SARS-CoV-2 biology, including whether it increases the risk of cancer. This study aims to identify the molecular interactions between COVID-19 infections and cancer processes.

**Materials and Methods:** We integrated recent data on SARS-CoV-2 – host protein interactions, risk factors for critical illness, known oncogenes, tumor suppressor genes and cancer drivers in EpiGraphDB, a database of disease biology and epidemiology. We used these data to reconstruct the network of molecular links between SARS-CoV-2 infections and cancer processes in various tissues expressing the angiotensin-converting enzyme 2 (ACE2) receptor. We applied community detection algorithms and Gene Set Enrichment Analysis (GSEA) to identify cancer-relevant pathways that may be perturbed by SARS-CoV-2 infection.

**Results:** In lung tissue, the results showed that 4 oncogenes are potentially targeted by SARS-CoV-2, and 92 oncogenes interact with other human genes targeted by SARS-CoV-2. We found evidence of potential SARS-CoV-2 interactions with Wnt and hippo signaling pathways, telomere maintenance, DNA replication, protein ubiquitination and mRNA splicing. Some of these pathways were potentially affected in multiple tissues.

**Conclusions:** The long-term implications of SARS-CoV-2 infection are still unknown, but our results point to the potential impact of infection on pathways relevant to cancer affecting cell proliferation, development and survival, favoring DNA degradation, preventing the repair of damaging events and impeding the translation of RNA into working proteins. This highlights the need for further research to investigate whether such effects are transient or longer lasting. Our results are openly available in the EpiGraphDB platform at https://epigraphdb.org/covid-cancer and the repository https://github.com/MRCIEU/covid-cancer (https://doi.org/10.5281/zenodo.6391588).

## Introduction

The coronavirus SARS-CoV-2 remains a major public health concern since it was first identified over 2 years ago, and a global scientific effort is focused on understanding its biology and how to minimize its clinical sequelae.

One major cause of concern is the virus’s unknown long-term effects [1]. It has long been known that viral infections have the potential to alter a cell’s DNA, activating carcinogenic processes and preventing the immune system from eliminating damaged cells. There is, therefore, an urgent need to understand the long-term health consequences of SARS-CoV-2, including whether it increases the risk of cancer or accelerates its progression [2].

A key to addressing these questions is to develop our understanding of the shared biology of COVID-19 disease and cancer, with the aim of identifying molecular mechanisms that may link SARS-CoV-2 with cancer. Early work from Gordon *et al*. [3,4] established a milestone in our understanding of the virus, describing viral bait genes and human target genes. Subsequently, Tutuncuoglu *et al*. [5] examined the structure and biological processes of this network of target genes, describing potentially undesirably perturbed candidate cancer genes at the host-virus interface. It has been hypothesized that SARS-CoV-2 may play a role in promoting cancer onset, for example by inhibiting tumor suppressor genes [6], and its potential impact on cancer risk and treatment [2]. However, to the best of our knowledge, the downstream links between SARS-CoV2 and established cancer genes have not been quantitatively described.

In this study, we hypothesize that these target genes may not be directly linked to cancer development themselves, but may interact with oncogenes, tumor suppressor genes and cancer drivers with the potential to influence oncogenic processes. To investigate these potential downstream effects, we analysed and visualized the network of interactions between the human target genes of SARS-CoV2 and cancer-related genes.

To develop this “COVID-cancer” interaction network we used the EpiGraphDB database, an analytical bioinformatics’ platform that supports population health data science, to link the recently mapped host-coronavirus protein interactions in SARS-CoV-2 infections with existing knowledge of cancer biology and epidemiology. Using these data, we constructed an accessible network of plausible molecular interactions between viral targets, genetic risk factors, and known oncogenes, tumour suppressor genes and cancer drivers for a variety of cancer types. Our results are openly accessible through the EpiGraphDB platform at https://epigraphdb.org/cancer-covid.

## Results

### Lung tissue

Coronaviruses target the respiratory system, and angiotensin-converting enzyme 2 (ACE2) produced by SARS-CoV-2 targets human cells expressing the ACE2 receptor [7], including in lung tissue. Thus, many patients with SARS-CoV-2 suffer long-term symptoms due to damage to the walls and lining of the lungs. We reconstructed the molecular network for lung cancer risk, representing cancer genes and interacting SARS-CoV-2 target genes (Figure 1). For the reconstructed network of other human cells expressing the ACE2 receptor see the interactive views at https://epigraphdb.org/cancer-covid or the data at https://github.com/MRCIEU/covid-cancer (https://doi.org/10.5281/zenodo.6391588).

**Figure 1.**
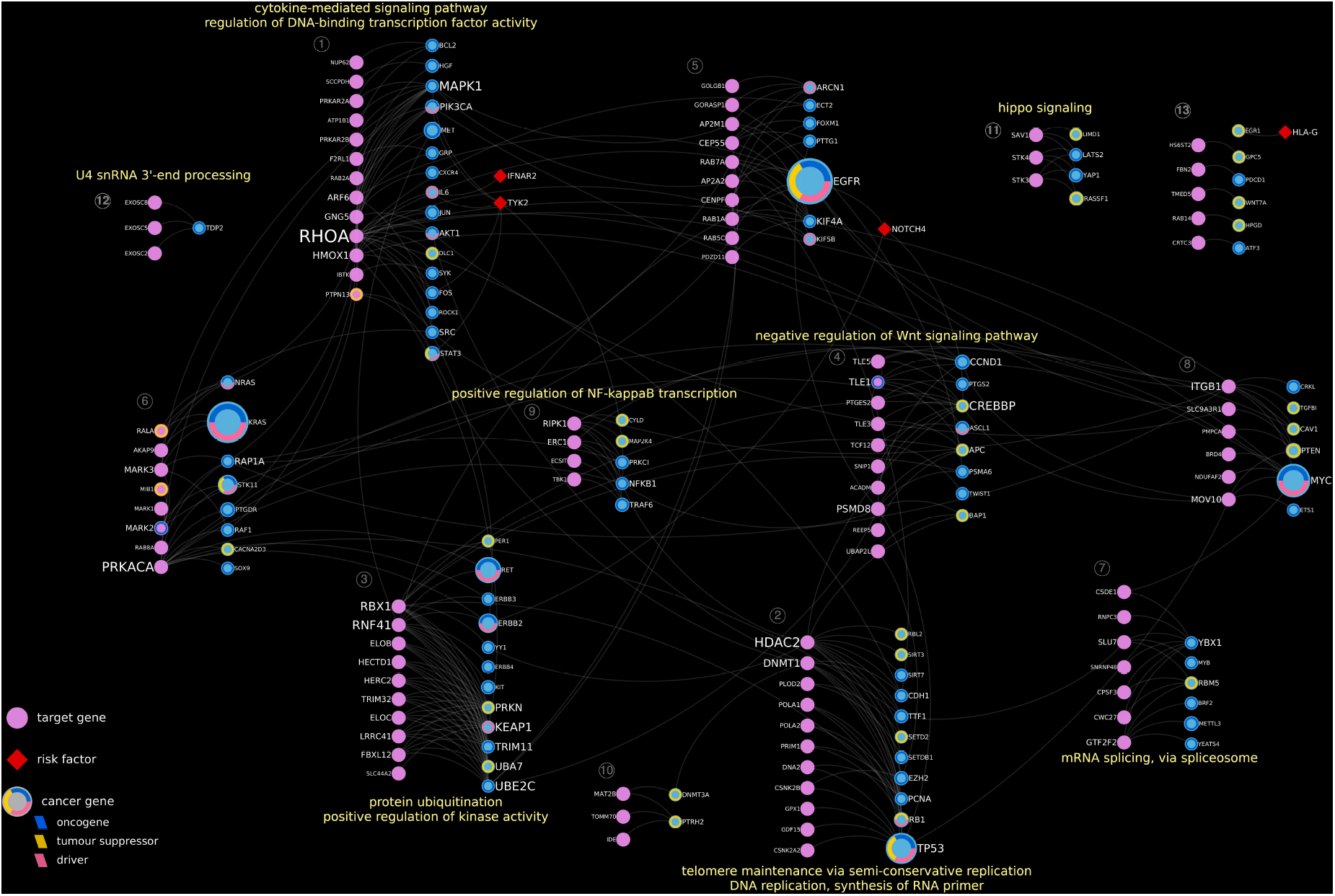
Network of protein-protein interactions between human genes targeted by SARS-CoV2 (pink), known carcinogenic genes in lung tissue (blue) and risk factors of critical illness (red), and selected enriched pathways for each cluster (indicated in grey). Node size is proportional to the node degree, and text size is proportional to the number of references in CancerMine [29] that cite each oncogenic gene, or the degree of the node in other target genes. The inner circle of each oncogenic gene node shows their role as oncogenes, tumor suppressor genes or cancer drivers.

We found 93 human genes targeted by SARS-CoV-2 which were either oncogenic (n=4) or interacted with oncogenic genes (n=92). These genes were clustered to enrich the network visualization, where each cluster of genes was depicted as two columns, one for SARS-CoV2 target genes and one for cancer genes. We then searched for molecular pathways that may be perturbed by each gene. A total of 12 clusters were found and 8 of them showed significant enrichment for pathways. The top enrichments in lung tissue are in Table 1, and the complete GSEA results for all tissues are in Supplementary File 1.

**Table 1.**
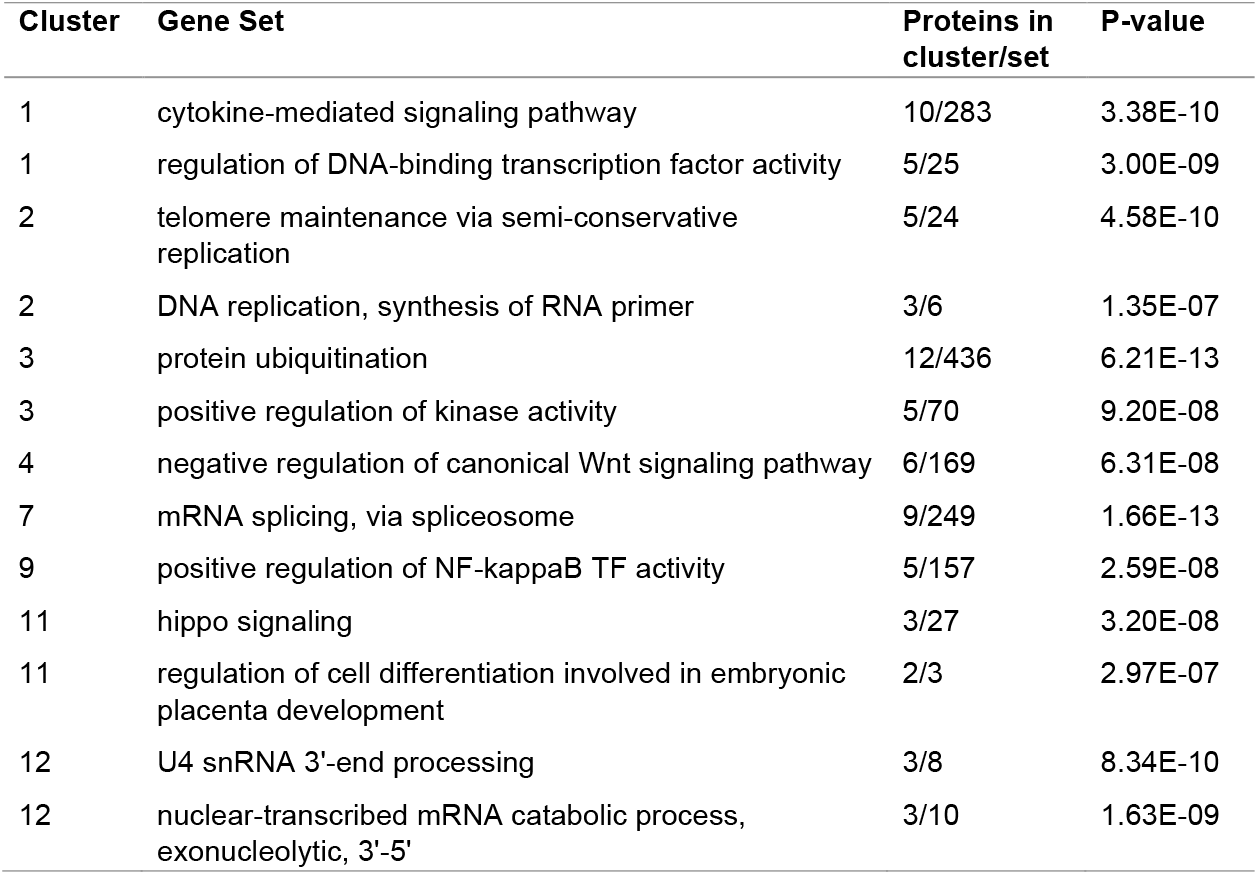
Gene set enrichment analysis (GSEA) for lung cancer. Only top two sets are shown per cluster. FDR threshold < 10^−4^.

| Cluster | Gene Set | Proteins in cluster/set | P-value |
| --- | --- | --- | --- |
| 1 | cytokine-mediated signaling pathway | 10/283 | 3.38E-10 |
| 1 | regulation of DNA-binding transcription factor activity | 5/25 | 3.00E-09 |
| 2 | telomere maintenance via semi-conservative replication | 5/24 | 4.58E-10 |
| 2 | DNA replication, synthesis of RNA primer | 3/6 | 1.35E-07 |
| 3 | protein ubiquitination | 12/436 | 6.21E-13 |
| 3 | positive regulation of kinase activity | 5/70 | 9.20E-08 |
| 4 | negative regulation of canonical Wnt signaling pathway | 6/169 | 6.31E-08 |
| 7 | mRNA splicing, via spliceosome | 9/249 | 1.66E-13 |
| 9 | positive regulation of NF-kappaB TF activity | 5/157 | 2.59E-08 |
| 11 | hippo signaling | 3/27 | 3.20E-08 |
| 11 | regulation of cell differentiation involved in embryonic placenta development | 2/3 | 2.97E-07 |
| 12 | U4 snRNA 3'-end processing | 3/8 | 8.34E-10 |
| 12 | nuclear-transcribed mRNA catabolic process, exonucleolytic, 3'-5' | 3/10 | 1.63E-09 |

Our results suggest SARS-CoV-2 host protein interactions could influence cancer risk through interactions with proteins in Wnt and hippo signaling pathways (clusters 4 and 11), two important pathways frequently linked to cancer due to their roles in cell proliferation, development and cell survival [8,9].

Perturbations in telomere maintenance and DNA replication may affect the integrity of DNA, favoring its degradation and preventing the repair of damaging events like gene fusions. The enrichment in cluster 2 suggests this could be influenced by SARS-CoV-2 infection. In addition, several target proteins interact with the quasi-universal tumor suppressor *TP53* [10] that regulates cycle arrest, apoptosis, senescence, DNA repair, or changes in metabolism.

SARS-CoV-2 infection may also impact on gene function through changes in the mRNA splicing process, impeding translation into working proteins, as depicted in cluster 7, and alterations in protein ubiquitination and the positive regulation of kinase activity as per the metabolic pathways enriched in cluster 3. In the latter, the oncogenes *RET* and *ERBB2* could be affected by upstream gene targets. RET (receptor tyrosine-protein kinase) is involved in numerous signaling pathways including cell proliferation and both cell death and survival [11]. In addition, ERBB2 is a protein tyrosine kinase that is part of several cell surface receptor complexes and member of the epidermal growth factor (EGF) receptor family that prevents the phosphorylation of the tumor suppressor gene *APC* [12]. Furthermore, *NOTCH4*, a gene increasing the risk of critical illness in people with COVID-19 [13], was linked to *CCND1* and *ERBB2*, genes that impact on cancer metastasis and poor prognosis [12,14,15].

Cluster 1 shows enrichment for the cytokine-mediated signaling pathway, including genes *IFNAR2* and *TYK2*. Both genes were identified as genetic risk factors for COVID-19 [13] and may interact with interleukin 6 (IL6), an important gene in the regulation of host defense during infections. Also, there was a significant enrichment of the regulation of DNA-binding transcription factors, including the oncogene *MAPK1* that plays a vital role in the regulation of meiosis, mitosis and postmitotic functions in differentiated cells [16].

### Cross-tissue comparison

We compared the oncogenic genes that are SARS-CoV-2 targets, or interact with them, in 6 different tissues expressing the ACE2 receptor, including lung, testis, colon, kidney, stomach and small intestine. Figure 2A shows the overlap of these sets of genes using a Venn diagram. Lung is the tissue with the most oncogenic genes (n=96) likely targeted and it shares 17 genes with colonic, 14 genes with gastric, and 6 genes with testicular, tissues. Of those, 13 genes are shared between lung, colonic and gastric, and 4 genes between lung, colonic and testicular, tissue.

**Figure 2.**
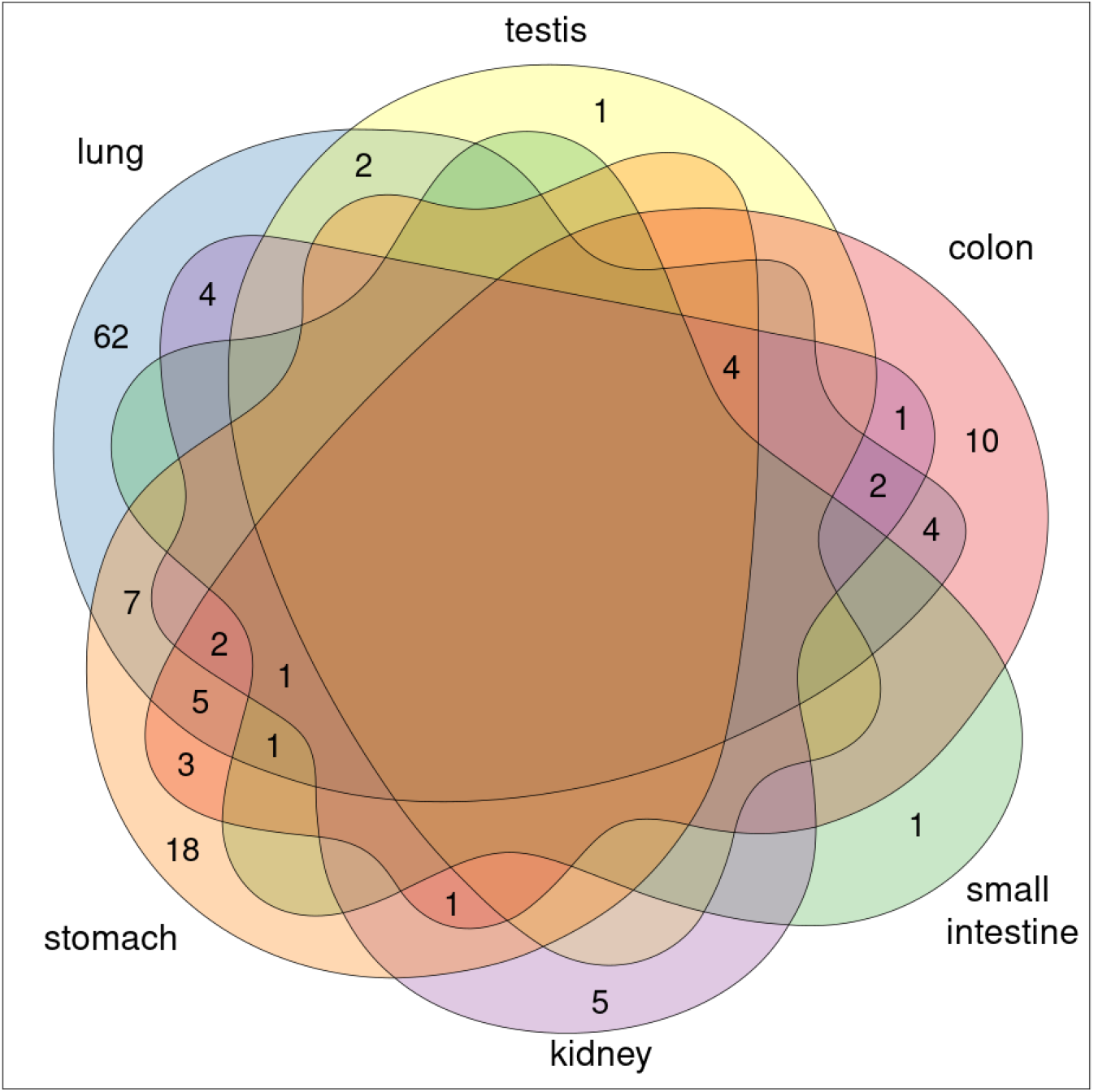
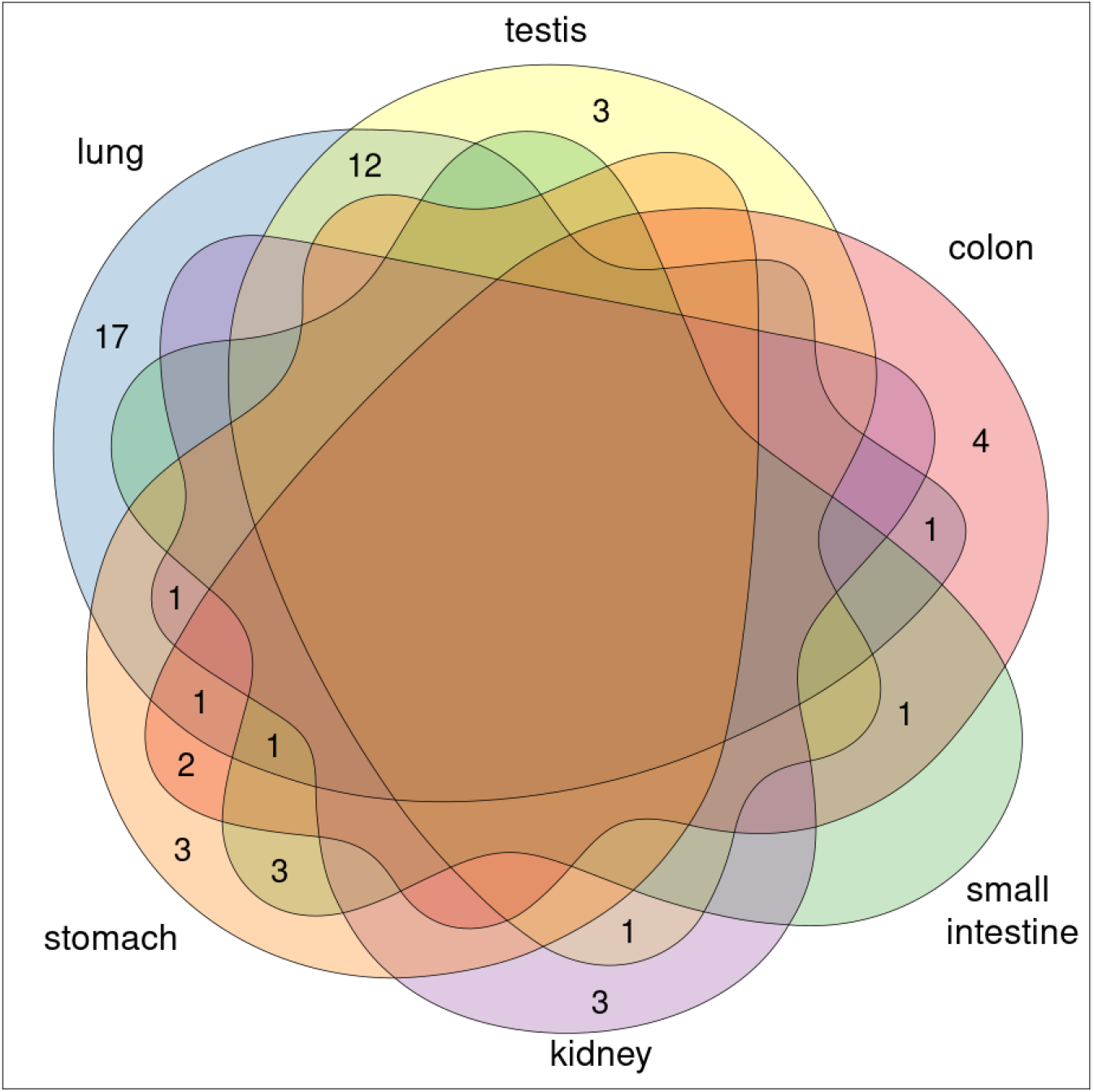
(A) Venn diagram with the number of oncogenic genes that are potentially perturbed in each of the tissues expressing ACE2 receptor. (B) Comparative results of the GSEA for each tissue, where each group depicts the number of enriched pathways.

Figure 3A details the genes shared between tissues. The well-established oncogenic genes *TP53, MYC, PTEN* and *RASSF1* were all present for kidney, stomach and small intestine. The oncogene *KRAS* is present in all tissue networks except testis and the tumor suppressor gene *APC* was shown to be targeted in lung, colon, small intestine and stomach.

**Figure 3.**
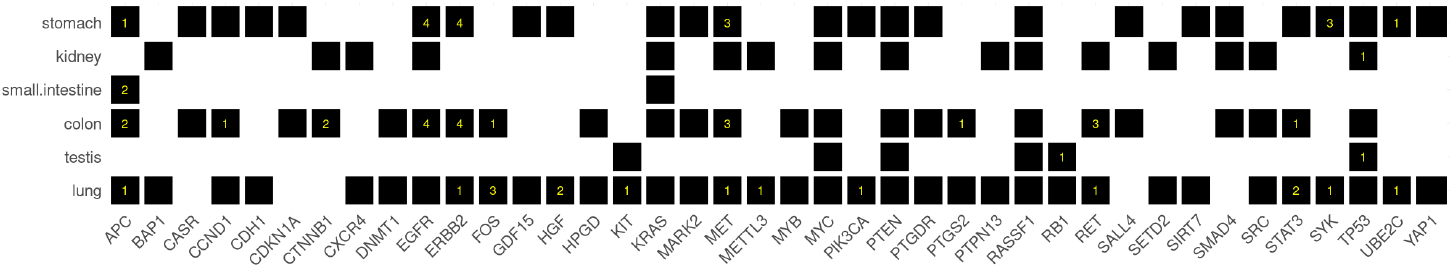
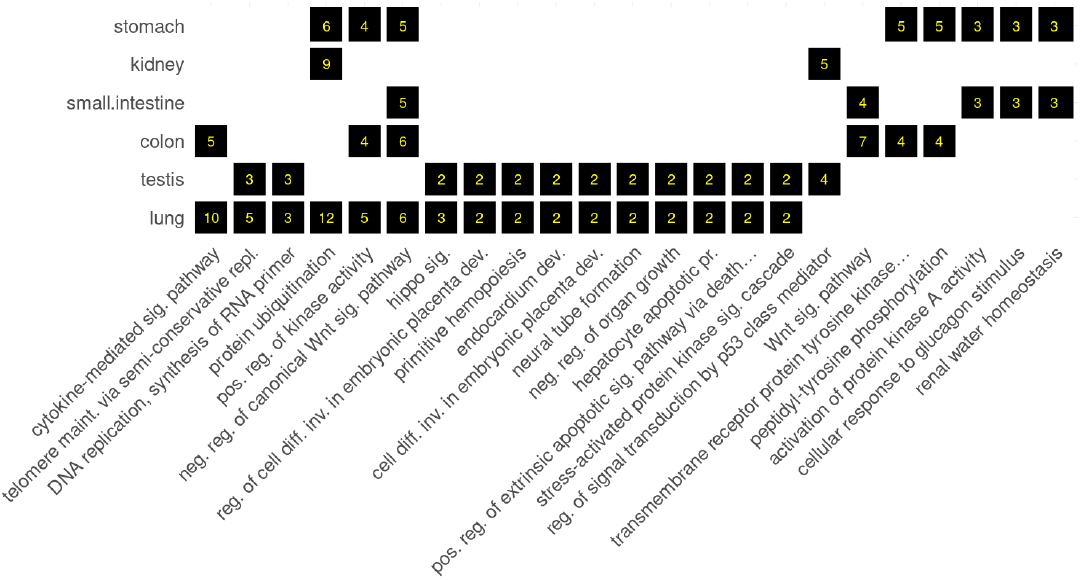
(A) Binary matrix representing the oncogenic genes that are potentially perturbed in each tissue. The numbers in yellow indicate in how many pathways each gene is part of the overrepresented set (FDR<10^−4^). Only those genes present in more than one tissue are depicted here. (B) Shared pathway enrichment between tissues, only showing pathways enriched in more than one tissue. Inside each box we indicated how many genes are overrepresented.

Looking at the molecular pathways (Figure 2B and 3B) there were 12 significantly enriched pathways shared between lung and testis. Those included important cancer-related molecular processes such as DNA replication, telomere maintenance, and the hippo signaling pathway. However, it is worth noting that this enrichment was triggered by only two genes *STK3* and *STK4*. STK3 and STK4 are key components of the Hippo signaling pathway, playing a pivotal role in tumor suppression by restricting proliferation and promoting apoptosis [17].

Moreover, the regulation of the canonical Wnt signaling pathway was enriched for lung, colon, small intestine, and stomach. Only testis showed enrichment for regulation of signal transduction by p53 class mediator. A detailed list of the genes and tissues in this comparative analysis between tissues can be found in Supplementary File 2.

## Discussion

It is estimated that 12% of all human cancers are caused by viruses. Viral oncogenesis is a complex process where viruses are necessary but not sufficient for cancer development and cancers only appear after long-term persistent infections [18]. Ghavasieh *et al*. [19] compared the host-parasite protein-protein interaction (PPI) network of SARS-CoV-2 against 92 known viruses finding similarities with respiratory viruses (Influenza A and Human Respiratory Syncytial virus), as would be expected, and also with other viruses such as HIV1, a risk factor for cancer due to its activation of chronic inflammation and the exhaustion of T cells.

Tutuncuoglu et al. [5] engineered the map of SARS-CoV-2 proteins directly interacting with potential cancer genes with the aim of identifying if coronaviruses and cancer cells disturb the same host pathways. Building on this approach, and with the objective of triangulating evidence [13], we extended the network of SARS-CoV-2 interacting proteins to include not just those which are oncogenic themselves (rarely the case), but also those cancer genes whose product interacts with target proteins (i.e. upstream or downstream of SARS-CoV-2 interacting proteins). Whilst the virus interacts with few oncogenes directly, some exceptions can be seen in Figure 1 for lung tissue. The tumor suppressor genes *PTPN13, RALA* and *MIB1* and the oncogene *MARK2* are direct targets of the virus and therefore more likely to be perturbed. The final functional impact is, however, not known.

Our results showed that SARS-CoV-2 could potentially alter fundamental cancer hallmarks [21]. Upstream PPI alterations in the Hippo signaling pathway, a key process in restraining cell proliferation and promoting apoptosis, may sustain a proliferative signaling and evasion of growth suppressors. The abnormal activation of the Wnt signaling pathway could result in a misregulation of the pluripotency and differentiation of cells, leading to the deterioration and metastasis of cancer. Furthermore, the alterations in the telomere maintenance pathway may enable the replicative immortality of cancer cells.

Understanding these enriched biological pathways may also help to explain the pathophysiology and differences in infection severity behind COVID-19. For instance, males have higher risk of severe SARS-CoV-2 infections [22], and the enrichment in testis for the signal transduction by p53 class mediator pathway (with the only other enrichment in kidney) may give us prospective clues about the mediators affecting the prognosis of COVID-19.

Hypothetically, an inhibited tumor suppressor such as p53 could have an oncogenic potential through mechanisms similar to HPV [6]. Nonetheless, well-known oncogenic viruses like HPV typically establish long-term infections, but there is no evidence of long-term viral latency of SARS-CoV-2 [6]. Exceptions to this are immunodeficient individuals, such as some cancer patients, who are at risk of rapid viral evolution and prolonged infection [23,24]. Long-term infections in these patients represent a source of variation for the virus that aids *intra*-host and *inter*-host transmission [25], and requires additional mitigation strategies such as self-isolation, immunization, combination monoclonal antibodies and antiviral agents. Similarly, the end of legal mitigation strategies may place some patients at risk of continuous reinfection and to a similar biological stress as in long-term viral infections.

To conclude, our novel results have pointed out the potential impact of SARS-CoV-2 on oncogenic pathways, and provide new understanding of the biology of SARS-CoV-2 virus and its molecular connection to cancer. However, the long-term implications of SARS-CoV-2 infection are still unknown and further research is needed to comprehend the oncogenic risk of SARS-CoV-2 and its induced inflammatory response.

## Materials and Methods

### Data sources and data integration

In this project we used the knowledge graph EpiGraphDB [26], available at https://epigraphdb.org, to extract the human interactome of protein-protein interactions (PPI) originally obtained from StringDB [27]. Only high confidence interactions (interaction score > 700) were included in the database.

We extended the dataset with the SARS-CoV-2 human protein–protein interactions that were extracted from the seminal work of Gordon *et al*. [3,4]. Only human target proteins were added to the network.

We also included the genetic risk factors of critical illness in COVID-19 identified by [13]. A total of 9 single nucleotide variants (SNPs) were mapped to the nearest gene. Those were related to key mediators of inflammatory organ damage and other host antiviral defense mechanisms.

Finally, a list of genes previously associated with tissue specific cancers was extracted from CancerMine (http://bionlp.bcgsc.ca/cancermine/) [29], a database based on automated mining of the scientific literature to identify and classify cancer genes as oncogenes, tumor suppressor genes and cancer drivers for different cancer types. Note that these are non-exclusive categories, and each gene can have multiple roles. Only genes with more than 2 references in the literature for any cancer type were included to avoid low confidence results.

When necessary, conversion of gene names to UniProt identifiers, or vice versa, was performed using BioMart [28].

### Network reconstruction

Network approaches have the potential to provide a holistic representation of unrelated genes or proteins onto a large-scale network context. Here, we built accessible networks of plausible molecular interactions between viral targets, genetic risk factors, and known oncogenes, tumor suppressor genes and cancer drivers for different cancer types.

It has been recently demonstrated that SARS-CoV-2 targets specific human cells that express the receptor ACE2 [7]. Hence, we reconstructed 6 different networks, one for each tissue where ACE2 is highly expressed: lung, testis, colon, kidney, stomach, and small intestine. Each one of these networks was constructed based on the SARS-CoV-2 human target proteins, tissue-specific cancer oncogenic genes that interact with these target proteins, and genetic risk factors of critical illness of COVID-19 that interact with any cancer gene.

These networks were compiled as CSV files and loaded into the Cytoscape platform, version 3.8.2 [30]. The scripts to reproduce the network construction are available at https://github.com/MRCIEU/covid-cancer (https://doi.org/10.5281/zenodo.6391588).

### Community detection and GSEA

To increase the statistical power in our protein-protein interaction networks we reduced the dimensionality of data using network modules and functional annotation. These techniques and their accompanying visualizations tackle the challenge of deriving relevant biological insights from long lists of genes and allow us to gain novel insights about the true biologically important effect, leading to a more intuitive interpretation of the data.

We used an edge-betweenness algorithm [31], as implemented in the Cytoscape plugin ReactomeFIPlugIn version 8.0.3 [32], to partition the networks into modules of interacting proteins. Only modules with more than 4 nodes were allowed.

Subsequently, we looked at the enrichment analysis for pathways and GO annotations for each subset of genes and annotated the modules, using the binomial test implemented in ReactomeFIPlugIn. Top enriched categories with a cutoff FDR<10^−4^ were extracted for each module.

All network diagrams were drawn with Cytoscape, and layouts were tuned manually to clearly display the modules and different target genes and cancer related genes.

### Venn diagrams and binary matrix

The comparison of targeted oncogenic genes and the pathway enrichment between different tissues were performed using Venn diagrams with the R package *venn* [33]. For a more detailed representation of similarities between tissues we converted the data into binary matrices for genes and pathways. These matrices were plotted with the R package ggcorrplot [34], but only those genes and pathways shared between two or more tissues were depicted. R version 4.1.2 [35].

## Author contributions

P.E. and T.R.G. wrote the manuscript. P.E. conceptualized the project. T.R.G. and P.E. supervised the project. P.E., T.R.G and R.M. acquired funding for the project. P.E. integrated the data and performed the data analysis. All authors contributed to the review and editing of the manuscript and provided comments.

